# Effects of ketamine on frontoparietal interactions during working memory in macaque monkeys

**DOI:** 10.1101/2023.05.16.540957

**Authors:** Liya Ma, Nupur Katyare, Kevin Johnston, Stefan Everling

## Abstract

Schizophrenia is known as a syndrome of dysconnection among brain regions. As a model for this syndrome, low doses of N-methyl-D-aspartate (NMDA) receptor antagonists, such as ketamine, produce schizophrenia-like symptoms and cognitive deficits in healthy humans and animals. One of such deficits is impaired working memory, a process that engages an extended network of both frontal and parietal areas. While ketamine is known to disrupt working memory by altering both spiking and oscillatory activities in the lateral prefrontal cortex (lPFC), it remains unknown whether NMDA receptor antagonists also produce frontoparietal dysconnection during working memory processes. Here, we simultaneously recorded both single unit activities and local field potentials from lPFC and posterior parietal cortex (PPC) in macaque monkeys during a rule-based working memory task. Like previous work in the lPFC alone, we found that ketamine compromised delay-period rule coding in single neurons and reduced low-frequency oscillations in the PPC. Furthermore, ketamine reduced task-related connectivity in both fronto-parietal and parieto-frontal directions. Consistent with this, ketamine also weakened interareal coherence between spiking and low-frequency oscillatory activities. Our findings demonstrate the utility of acute NMDA receptor antagonist in simulating a syndrome of dysconnection and support this model in its potential for the exploration of novel treatment strategies for schizophrenia.

## INTRODUCTION

Working memory is at the foundation to many of our daily tasks, and its impairment is a transdiagnostic symptom in neuropsychiatric disorders such as schizophrenia [1,2]. While the positive symptoms in schizophrenia are often well-controlled with antipsychotics, currently there is no effective treatment for the impairment of working memory, among other cognitive deficits [3,4]. Working memory is known to engage an extended brain network which includes the lateral prefrontal cortex (lPFC) and the posterior parietal cortex (PPC) [5–7]. Meanwhile, structural and functional dysconnection has been regarded as a key feature of schizophrenia [8–10]. Therefore, a viable model for working memory impairment in the syndrome should provide insight into the role played by intra-and inter-areal communication in the process.

The N-methyl-D-aspartate receptor (NMDAR) glutamate receptor plays an important role in working memory [11,12]. In healthy human participants, a low dose of Ketamine (at around 0.4mg/Kg), an NMDA-receptor antagonist, is known to produce short-lasting positive and negative symptoms as well as cognitive deficits, resembling those observed in people with schizophrenia [13–15]. In both experimental rodents [16](Homayoun et al., 2004, Condy et al., 2005, Enomoto and Floresco, 2009) and non-human primates [17–19], a similarly low dose of ketamine led to cognitive impairment comparable to those of ketamine-treated healthy humans and patients with schizophrenia. Previously, using single-unit recordings in the lPFC of macaque monkeys, we revealed that ketamine weakened working-memory related signals and increased trial-to-trial variability at the levels of single neurons and neuronal ensembles [20]. We also found that ketamine increased gamma-band activity (>30Hz) while decreasing the beta (13-30Hz) rhythm in lPFC local field potentials (LFPs), which was significantly correlated with working-memory performance [21].

Neurons in the PPC are known to respond during working memory in similar ways as those in the lPFC [22–30], despite differences in their intrinsic properties [31–34]. This observation points toward a role of real-time frontoparietal interaction in working memory performance. Evidence of frontoparietal dysconnection has been reported in several studies with schizophrenia patients [35–37] and was correlated with impaired working memory performance [38–40]. Specifically, NMDA receptor antagonism is known to profoundly change the cell-and network-level dynamics in frontal and parietal cortices of macaque monkeys, during a cognitive control task [41]. However, the physiological mechanisms underlying this functional dysconnectivity is yet to be fully characterized.

To understand the network-level mechanism underlying the working memory deficit following ketamine treatment, we recorded single-unit activities and LFPs from both lPFC and PPC in macaque monkeys performing a working memory task [17](Figure 1A). In each trial, a color cue was presented which mapped onto the pro-or anti-saccade rule in working memory. Among these rule visible trials (RS), we interleaved rule memorized trials (RM) in which a delay was inserted between the cues and target onset, when the monkeys fixated on a white dot. Given our previous findings [17,20], we expected to find a weakening in rule-coding among PPC neurons. Additionally, we expected similar bidirectional changes in gamma and low-frequency oscillatory power in the PPC as in the lPFC [21]. Importantly, we hypothesized that NMDA antagonism would result in dysconnectivity in the frontoparietal network, in a way related to working-memory performance.

**Fig 1.**
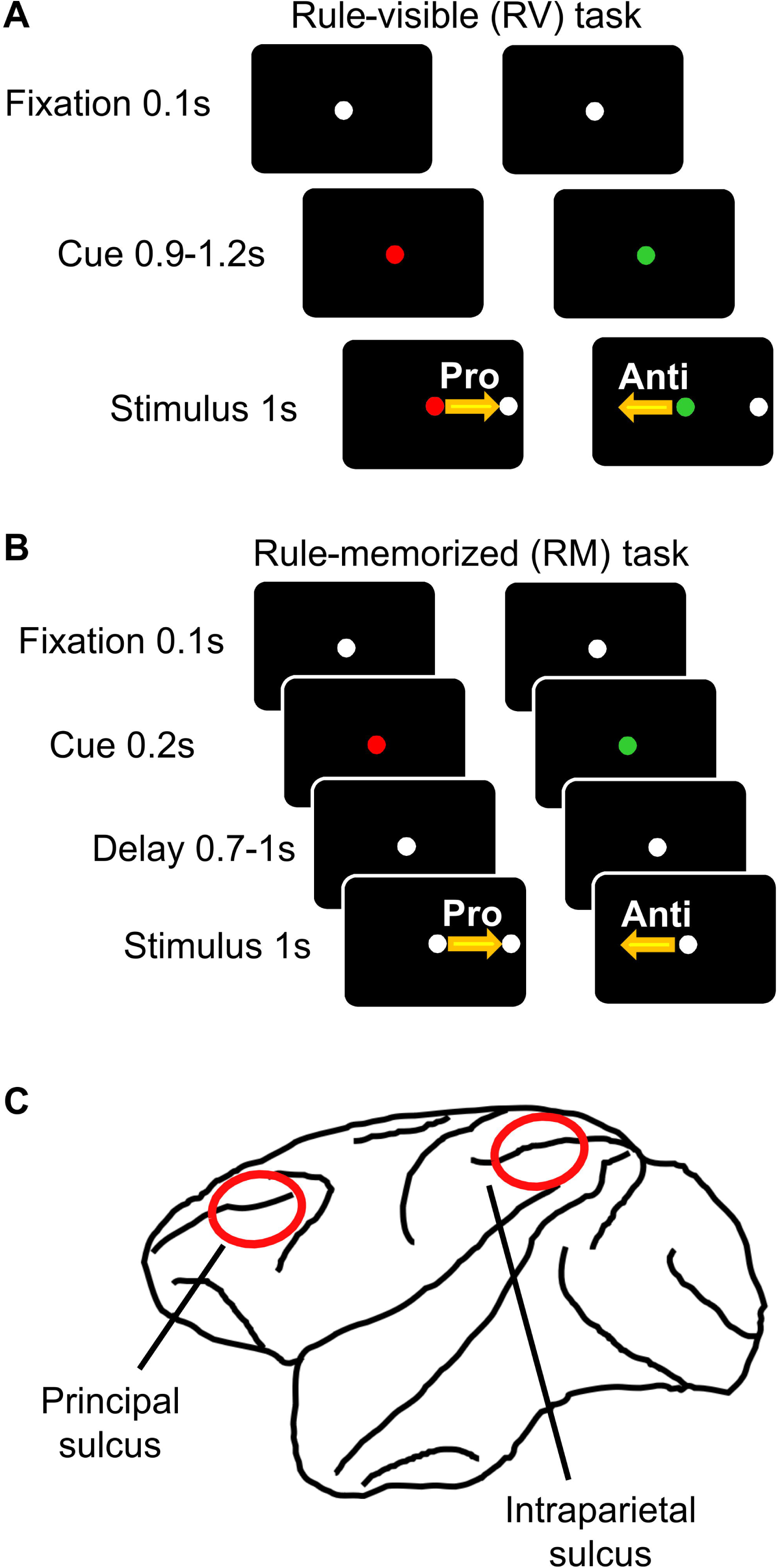
Schematics for the design of the rule-visible (**A**) and the rule-memorized (**B**) pro-and anti-saccade task, and the placement of recording chambers (**C**).

## RESULTS

On average, monkeys completed 528 trials (excluding omitted or no-response trials), including randomly interleaved pro-and antisaccade trials in the RV and RM conditions, per session, with sessionsranging from 41 to 65.5 minutes in duration. Ketamine had a significant effect on percentage correct responses compared to saline administration (mixed-model ANOVA, main effect of drug: *F*_1,156_ = 5.4, p = 0.024). Following ketamine injection, performance deteriorated in the first 10-min interval in trials under both rules and in both tasks (post hoc Tukey’s test, p <= 0.00011; Figure 2A). By the 3^rd^ 10-min interval, performance in each trial type showed recovery in comparison to either the 1^st^ or the 2^nd^ 10-min interval after ketamine injection (p <= 0.01). Notably, both RV and RM antisaccade trials were performed more poorly than prosaccades (p <= 0.00070; Figure 2A, black vs gray lines) in the 2^nd^ 10-min post-injection interval, even though pro-and anti-saccades were performed with equal accuracy before ketamine injections (p >= 0.84). Thus, consistent with previous reports [17,20,21], ketamine had a stronger impact on antisaccade performance. By contrast, the performance across tasks and rules did not change following a saline injection (p > 0.99; Figure 2B).

**Fig 2.**
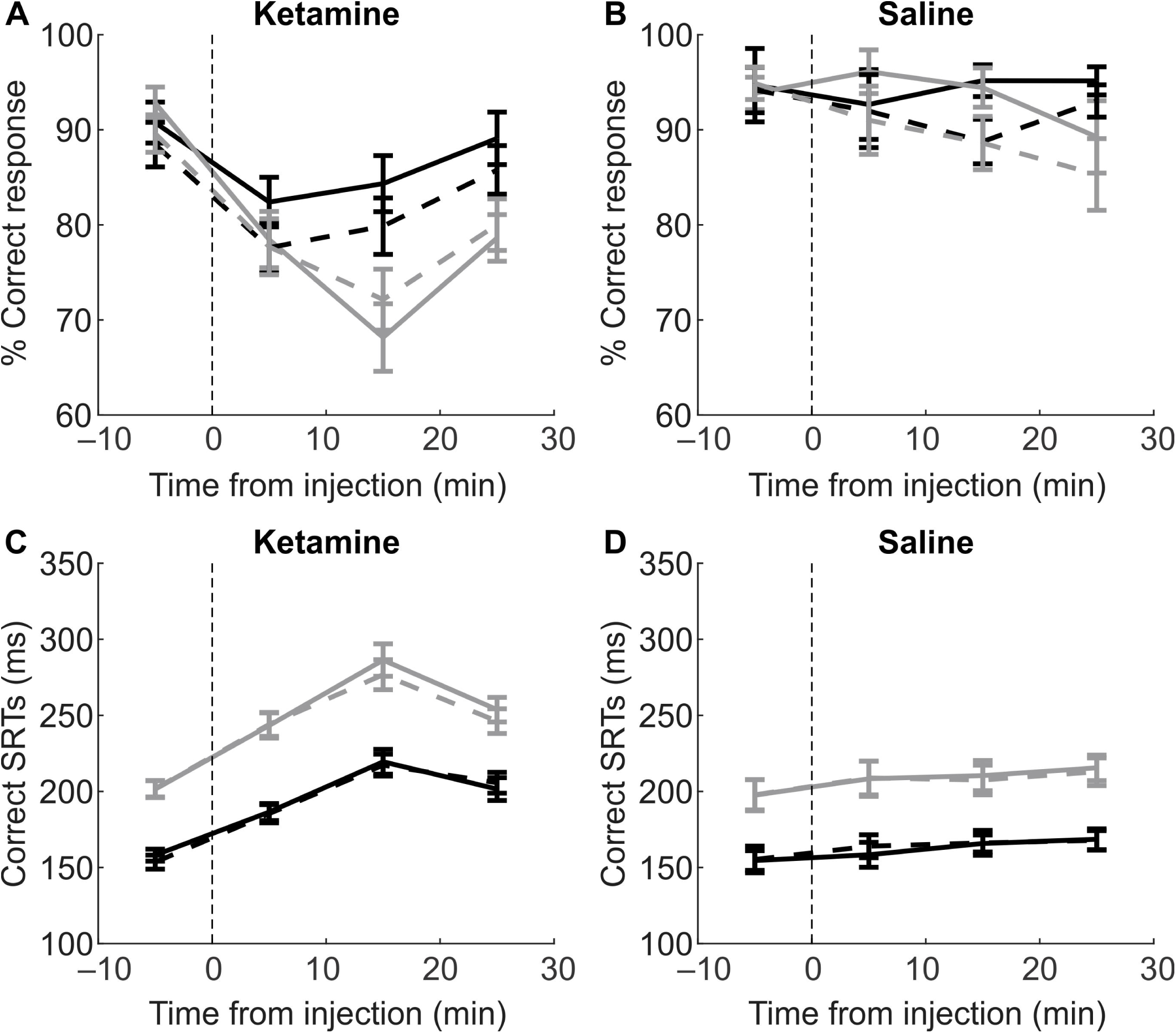
Effects of ketamine **(A, C)** and saline **(B, D)** injection on the percentage of correct response **(A, B)** and saccadic reaction times **(C, D)**. Performance on RV trials are shown as solid lines, and RM trials as dashed lines. Prosaccade trials are shown in black, antisaccade trials in gray. Error bars indicate the standard deviation of the mean.

Furthermore, ketamine, but not saline, significantly increased the saccadic reaction times (SRTs) (mixed-model ANOVA, effect of drug, *F*_1,150_ = 6.26, p = 0.016; Figure 2C, D). This was true for all trial types (post hoc Tukey’s test: ketamine: p = 2.8 × 10^-5^, saline: p >=0.98). While antisaccades had longer SRTs (effect of rule: *F*_1,150_ = 118.82, p = 8.1 × 10^-15^; gray curves), the effect of ketamine on these was equivalent to that on prosaccades (no drug-rule interaction: *F*_1,150_ = 1.23, p = 0.27). The monkeys’ performance was also similar across RV and RM trials (no task effect: *F*_1,150_ = 1.65, p = 0.21; solid vs dashed lines).

### Ketamine altered LFP powers in both lPFC and PPC

In 45 sessions involving ketamine injections, we recorded from a total of 152 and 171 channels in the lPFC and PPC, respectively, from which we isolated 262 and 248 single units. Here we consider only electrodes from which well-isolated units were detected. As previously reported [21], ketamine significantly weakened low-frequency LFP oscillations in the lPFC (blue with black contours, Figure 3A) in both inter-trial intervals (ITIs) and delay periods under both rules and in both tasks. While the increase in gamma rhythm did not reach significance (orange and red areas), the direction of this change is consistent with our previous report. Additionally, we observed the same bidirectional change in the PPC (Figure 3B).

**Figure 3.**
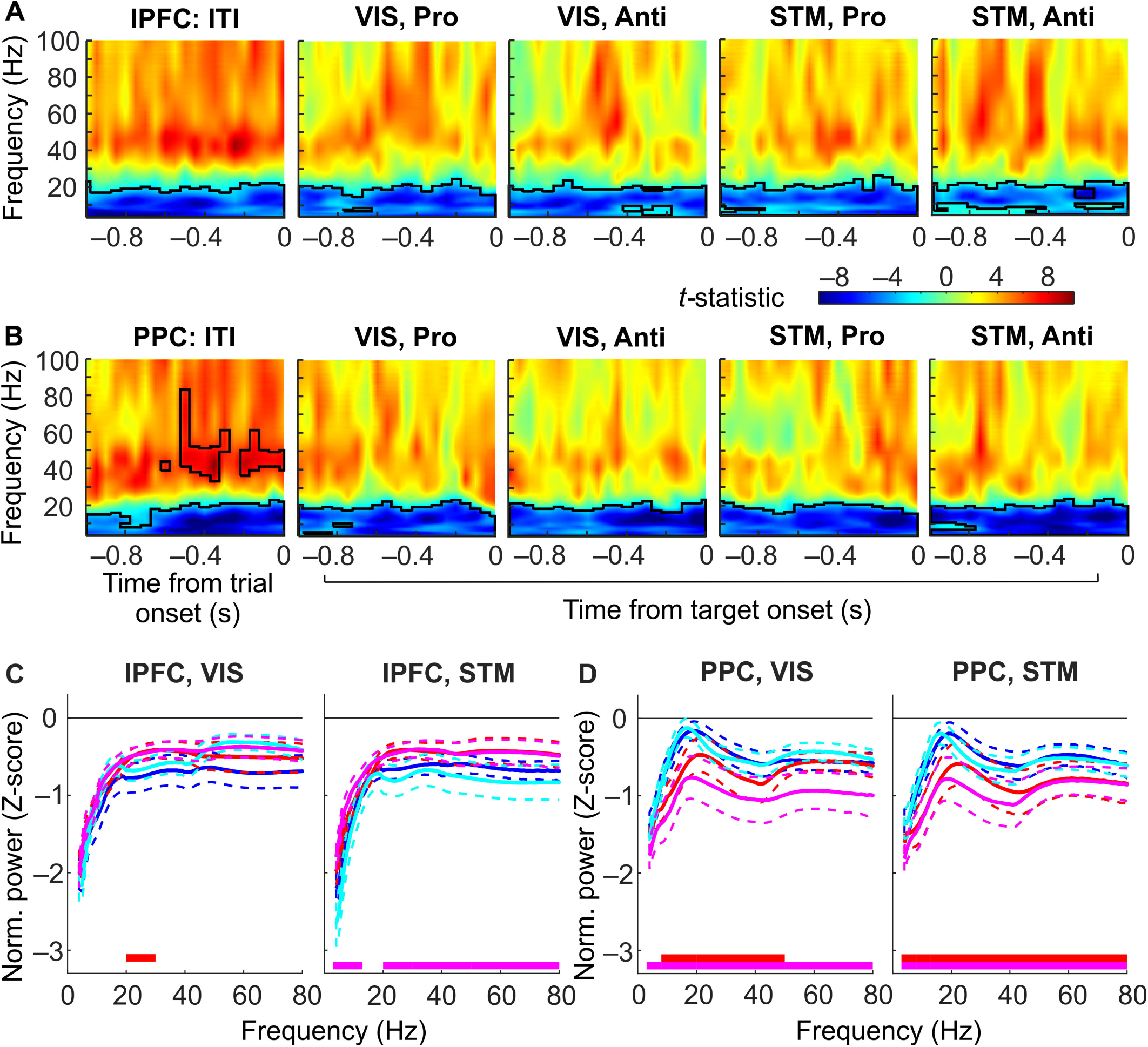
The effect of ketamine on LFP power in lPFC **(A)** and PPC **(B)**, and on task-specific oscillatory power **(C and D)** during RV (left panels) and RM (right panels) tasks in the two areas. A and B shows t-statistics between pre-and post-injection LFP power spectra. During both the ITI (left panel) and delay period (middle and right panels), and across both areas, low frequency oscillations in the theta, alpha and beta bands weakened (cool colors) after ketamine injection. By contrast, gamma-band power tended to increase (warm colors), especially during the ITI. Black contours indicate significant changes by permutation test (p<0.01). C,D show delay-period LFP power normalized against ITI. **C)** In the lPFC, ketamine increased this task-related LFP in high beta band in RV prosaccade trials, and in all bands except low beta in RM antisaccade trials. **D),** In the PPC, ketamine reduced task-related LFP power in several bands across tasks and rules in the PPC. Horizontal red (prosaccades) and magenta (antisaccades) bars indicate frequency bands with p<0.05 in repeated-measures ANOVA.

Going beyond the general effect of ketamine, to quantify any task-related effect of ketamine, we decibel normalized the LFP power in delay periods against that in ITIs (Herrmann et al., 2014). Overall, ketamine had somewhat opposite effects on task-specific LFP power in the two areas: in the lPFC it *increased* the task-related LFP power (repeated-measures ANOVA, drug-task interaction: *F*_1,755_ = 10.63, p = 0.0014, Figure 3C), particularly in RV prosaccade (high beta, p = 0.00099) and RM antisaccade trials (theta, alpha, high beta and both gamma bands (p <= 0.0021, Figure 3C,D). In the PPC, it *decreased* the power in both RV (drug-task interaction: *F*_1,850_ = 6.84, p = 0.0097, Figure 3D; antisaccade: p = 5.0 × 10^-5^ in all bands; prosaccade: p <= 0.0030 in alpha, beta and low gamma) and RM tasks (p <= 5.2 × 10^-5^ in all frequencies, Figure 3E,F).

### Ketamine reduces coherence between lPFC and PPC in a task-dependent manner

We then asked if ketamine altered the interactions between lPFC and PPC, both non-directionally as in coherence and directionally as in Granger’s prediction. For coherence, we calculated the de-biased weighted phase lag index (dWPLI) [43].

We first examined whether ketamine affected coherence during the pre-trial ITIs. We found a significant drug-frequency interaction within lPFC (rmANOVA, F_5,1030_ = 4.2, p = 0.00089), within PPC (F_5,1350_ = 3.5, p = 0.0035) as well as across the two areas (F_5,2960_ = 4.8, p = 0.0002). Specifically, within each area, there was a reduction in theta-band coherence (post hoc Tukey test, p = 2.4 and 6.2 × 10^-5^ respectively). Additionally, ketamine reduced alpha-band coherence between lPFC and PPC (p = 2.2 × 10^-5^). These effects were small (eta-squared from 0.008 to 0.02) and were not observed in any other frequency band. Thus, ketamine weakened frontoparietal coherence during the off-task period in the lowest-frequency bands.

Next, to examine task-related coherence, we normalized the delay-period coherence against those calculated from the ITIs. In the lPFC, ketamine did not affect task-related coherence during either task (Figure 4A,B). By contrast, ketamine significantly reduced within-PPC coherence (repeated-measures ANOVA and post-hoc Tukey’s test, p<0.05, Figure 4C, D), especially in theta band during RM antisaccade trials (Figure 4E) and in low-beta band during RVantisaccade and RM prosaccade trials (Figure 4F).

**Figure 4.**
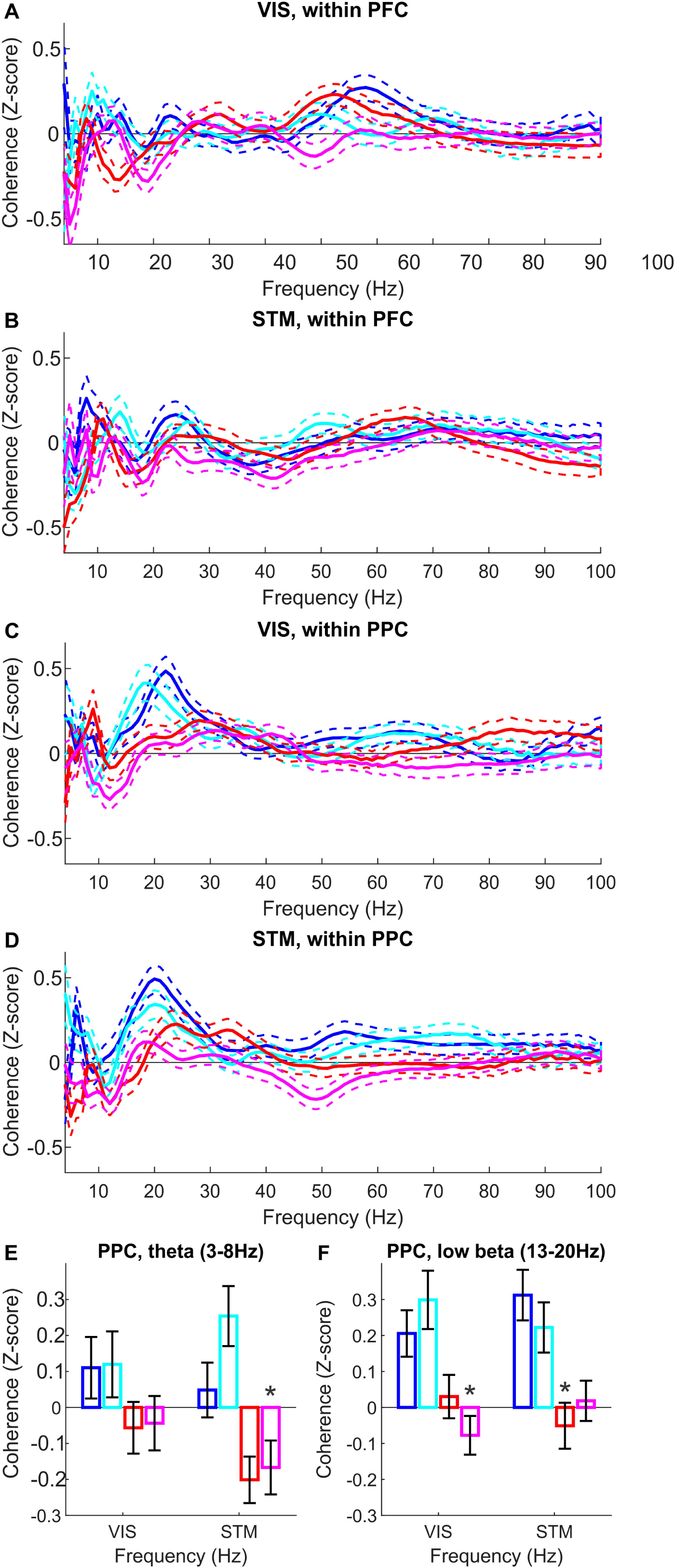
Effects of ketamine on coherence within lPFC and PPC. **A)** and **B)** On intra-lPFC coherence, ketamine had no significant effect in either task. **C)** Ketamine reduced intra-PPC coherence, especially in theta band in RM antisaccade trials, and **D)** in low-beta band in RV antisaccade and RM prosaccade trials.

We then examined lPFC-PPC task-related coherence, in which the effect of ketamine was found across even more frequencies (Figure 5A,B). For RV prosaccade trials, ketamine weakened coherence in theta and alpha band (repeated-measures ANOVA and post-hoc Tukey’s test, p<0.05, Figure 5C). For RM prosaccade trials, ketamine reduced interareal coherence in low-beta band. For RM antisaccade trials, this effect was found in the high-gamma band. In the high-beta band, ketamine eliminated the rule-related difference in the RV task. Thus, ketamine compromised within-and cross-areal non-directional connectivity in a task-dependent manner.

**Figure 5.**
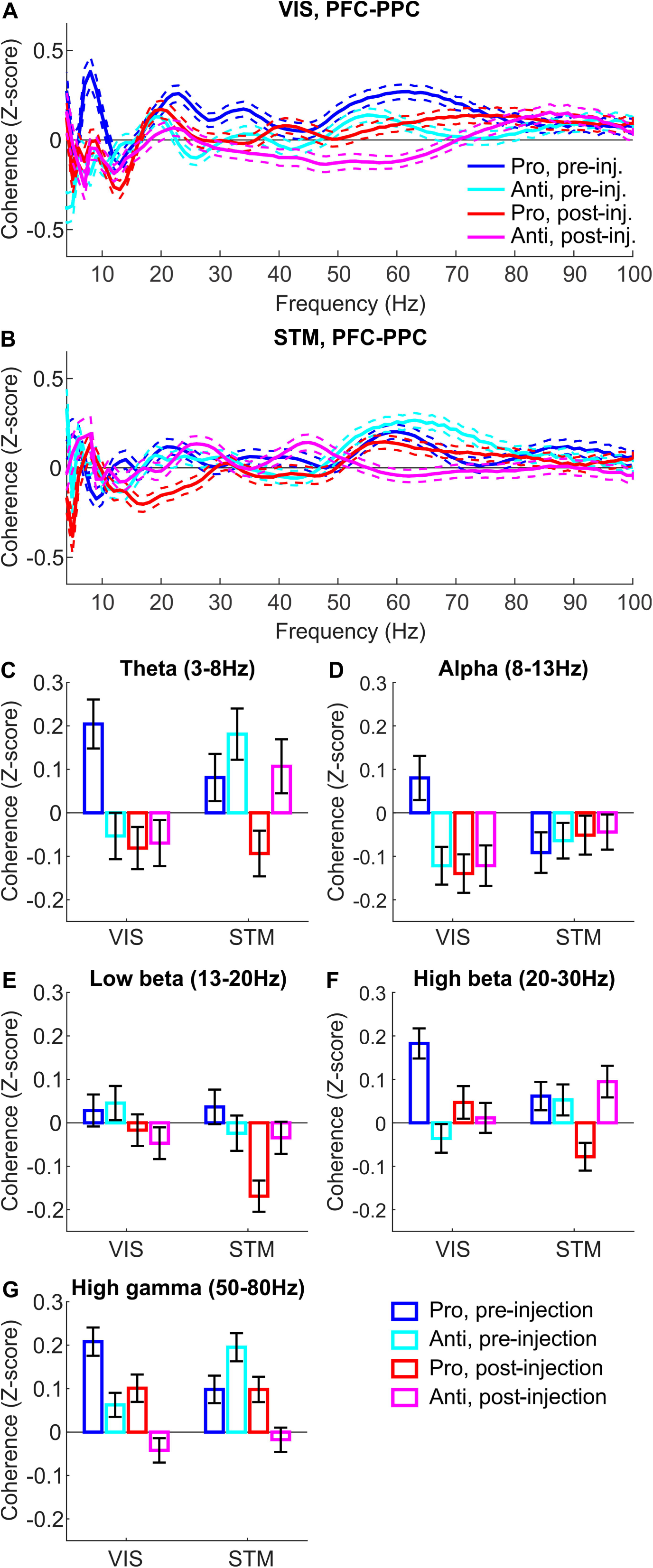
Ketamine affected lPFC-PPC coherence in multiple frequency bands in both **A)** RV and **B)** RM conditions. Ketamine significantly reduced coherence in RV prosaccade trials in **C)** theta and **D)** alpha band, in RM prosaccade trials in **E)** low beta band, and in RM antisaccade trials in **G)** high gamma band. In **F)** high-beta coherence, ketamine abolished the rule difference in the RV condition.

### Ketamine weakened lPFC-PPC and PPC-lPFC directional connectivity in the time domain

Results from the coherence analysis beget the question of whether the task-related connectivity was directional, and how ketamine might affect it. We therefore calculated time-domain Granger predictions [44] with lPFC leading PPC and vice versa. Before ketamine treatment, delay periods had lower connectivity than ITIs in both directions (rmANOVA and post hoc Tukey test, p = 3.2 × 10^-5^; Figure 6A, left). Additionally, during ITIs, the connectivity was equal across directions (p = 0.49; Figure 6A, left, empty bars), yet during the delay period, lPFC-PPC prediction became significantly stronger than PPC-lPFC prediction (p = 3.2 × 10^-5^; filled bars). Thus, the task modulated both the direction and strength of frontoparietal connectivity. After ketamine injections, connectivity weakened in both directions, during both delay periods and ITIs (p < 0.0001; Figure 6A, left vs right). Additionally, ketamine abolished the task-related modulation in both directions (p > 0.94; filled vs empty bars).

**Figure 6.**
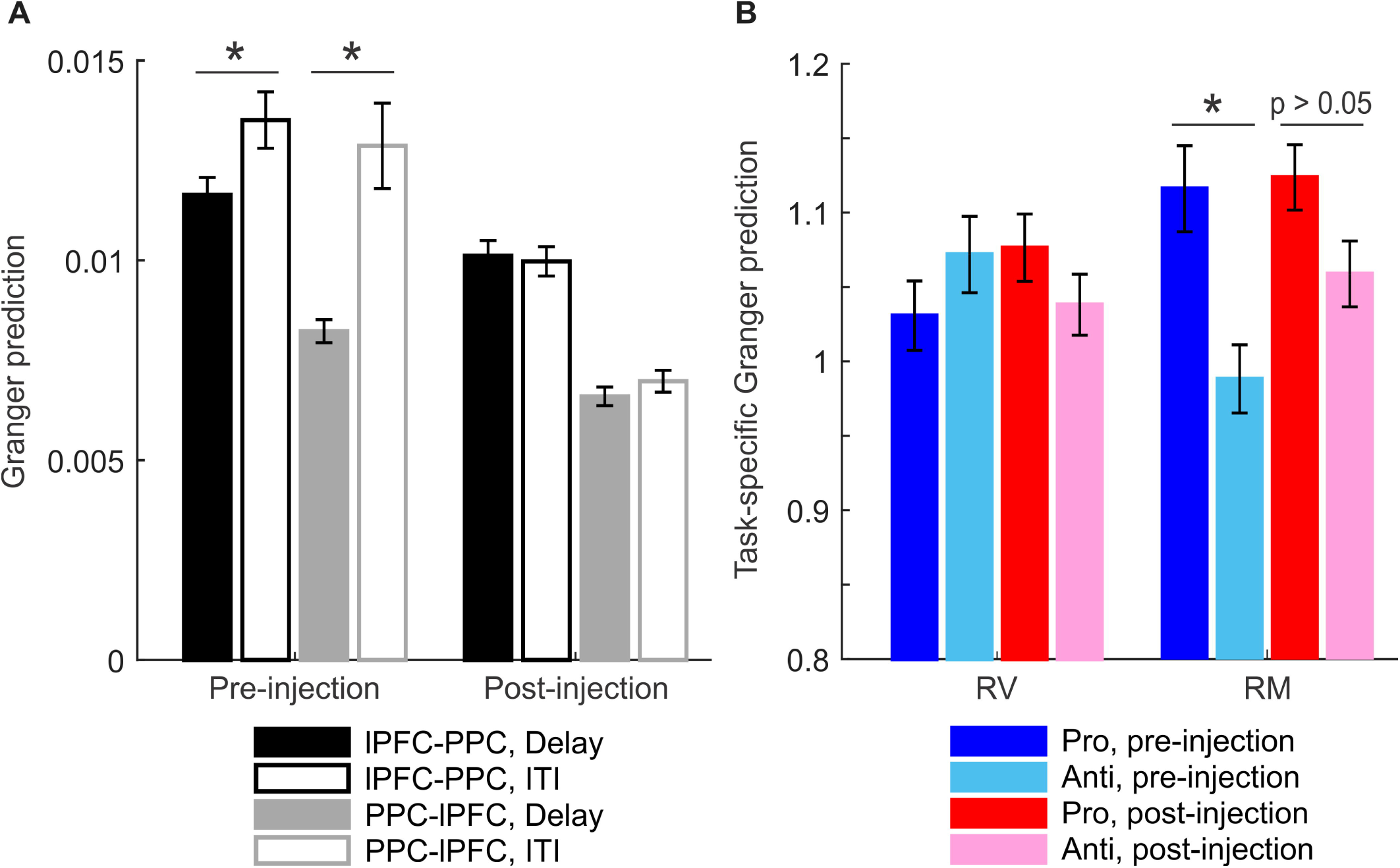
Ketamine affected the directional connectivity between lPFC and PPC in the time domain (Granger prediction). **A).** Granger prediction is overall stronger in the lPFC-lead than the PPC-lead model (left vs right set of bars). Ketamine significantly reduced their connectivity in both directions (black for pre-injection, gray for post-injection), and during both delay periods (filled bars) and inter-trial intervals (empty bars). **B).** Delay-period lPFC-lead Granger values were divided by ITI values to produce task-related directional connectivity. Ketamine did not affect task-related lPFC-PPC prediction in the RV task (left bars) but altered increased the values in RM trials (right bars: light blue vs pink). As a result, the difference between pro-and antisaccade trials (dark vs light blue bars) was reduced after ketamine injection (red vs pink bars).

To examine in greater detail any task-related effect of ketamine, we normalized the delay-period against the ITI Granger predictions. In the RV task, the lPFC-led connectivity did not differ by rule and ketamine did not affect the Granger values (Figure 6B, left bars). In the RM task, lPFC-PPC Granger values were greater during the prosaccade than antisaccade trials (p = 2.9 × 10^-5^); Figure 6B, blue bars, right side), and this difference was attenuated by ketamine (p = 0.054, red bars, right side). Additionally, the PPC-led connectivity showed an effect of rule in both RV and RM tasks, stronger in RV antisaccades and RM prosaccades (p < 6.3 × 10^-4^; Figure 6C, blue bars, both sides). Ketamine abolished both effects (p > 0.35, red bars, both sides). Taken together, ketamine not only affected frontoparietal connectivity in both directions, but also compromised rule-related modulations in this connectivity.

### Ketamine has task-specific effects on rule coding by lPFC and PPC neurons

After characterizing the effects of ketamine on frontoparietal field potentials, we went on to quantify its effects on information coding in spiking activities. We recorded a total of 262 and 248 neurons from the lPFC and PPC, respectively, with a mean pre-injection firing rate of 3.02 ± 5.89Hz (standard deviation) and 2.41 ± 3.20Hz. We conducted a mixed model repeated-measures ANOVA on the firing rates in four 10-minute intervals before and after ketamine administration. There was a significant effect of the drug (*F*_3,1524_ = 56.15, p < 1 × 10^-324^) and no effect of area (*F*_1,508_ = 2.00, p = 0.16). Consistent with our previous report [20], ketamine increased neuronal activities in the lPFC marginally in the first 10 minutes (p = 0.055) and significantly in the second and third 10-minute periods (p = 0.000032). In the PPC, its effect started in the first 10 minutes (p = 0.0015) and remained so in the later intervals (p = 0.00032). Additionally, in both areas, ketamine increased the coefficient of variation (CV) in inter-spike intervals, which reflects the irregularity in spiking activities (*F*_3,1518_ = 31.48, p < 1 × 10^-324^). Thus, ketamine increased the level as well as the irregularity in neuronal activities in the frontoparietal network.

We then quantified neuronal coding of the task rule by computing the d’ between the rules during fixation, cue presentation, delay and peri-saccadic periods for single units recorded from lPFC and PPC (see Materials and Methods; Figure 7A and B). Since RV trials had no delay period, the later portion of the cue period, which equaled to the delay period in duration, was used instead. Among lPFC units (Figure 7A), there was a significant rise in rule sensitivity from the cue to the cue/delay period (rmANOVA, effect of epoch: *F*_3,2955_ = 60.8, p < 1 x 10^-324^) in both tasks (post hoc Tukey’s test, RV: p = 0.002, RM: p = 0.008; solid and dashed black lines), which was abolished by ketamine administration (p = 0.99 and 0.58, gray lines). Cue/delay-period rule sensitivity was significantly reduced by ketamine in the RM (p = 0.02) and marginally in the RV task (p = 0.07). Among PPC units (Figure 7B), this rise in rule sensitivity was seen in the RV but not RM task (effect of epoch: *F*_3,2820_ = 30.7, p < 1 x 10^-324^, post hoc tests: RV: p = 0.0006, RM: p > 0.99; black lines), which also disappeared following ketamine injections (p > 0.99, gray solid line). Notably, the effect of ketamine on rule sensitivity was specific to the cue/delay period and was not observed in any other epoch.

**Fig 7.**
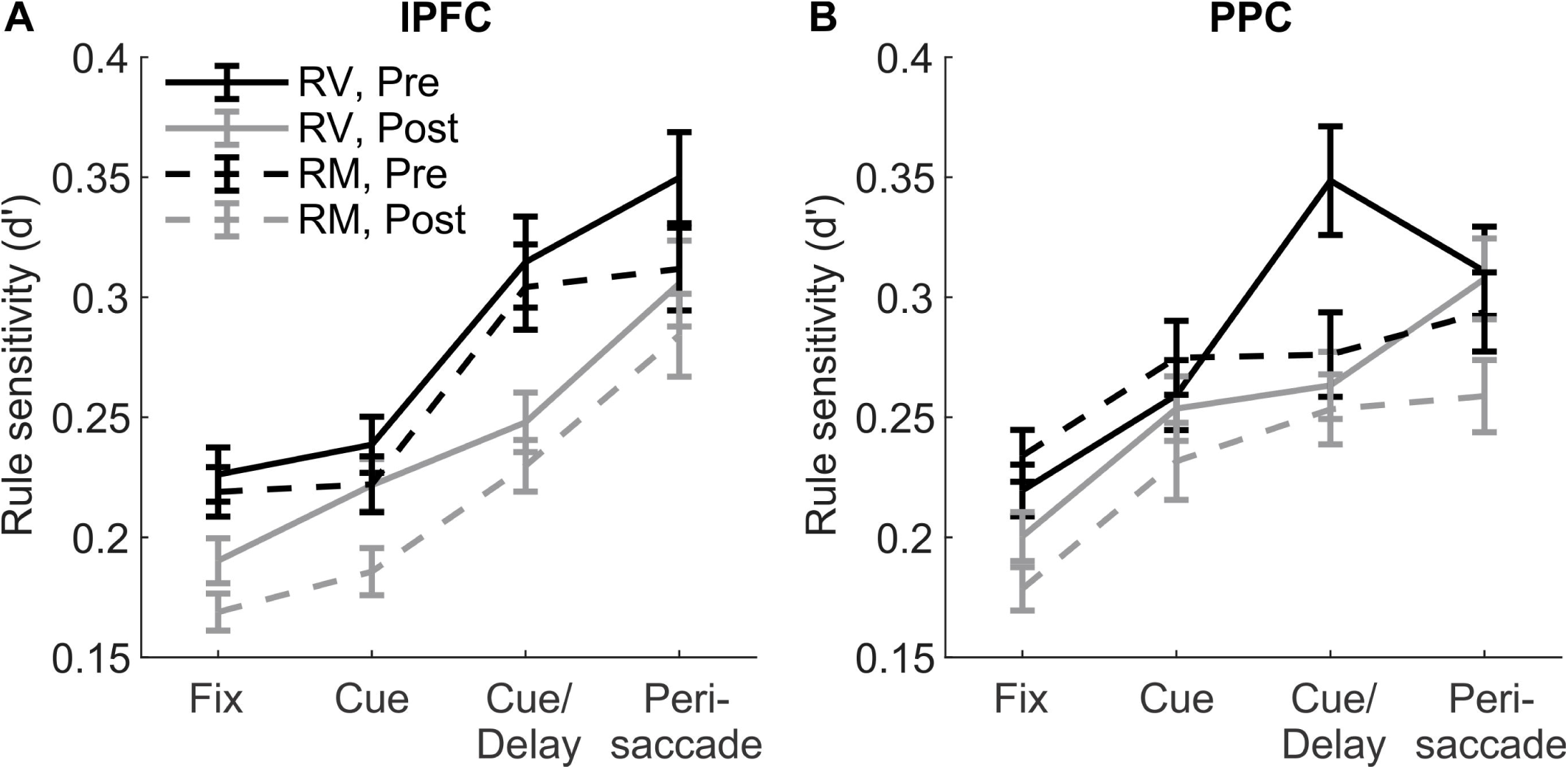
Rule sensitivity during different epochs in all neurons recorded from **A)** lPFC and **B)** PPC. The third epoch from VIS trials is the later part of the prolonged cue period, which matched the delay period in RM trials. Solid lines: VIS trials, dashed lines: RM trials. Pre-injection: black, post-injection: gray. Error bars indicate the standard deviation of the mean.

In both tasks, the red/green cues must be mapped onto pro-or anti-saccade rules based on memory, before the monkeys could choose the appropriate response. It is possible that this ‘mapping process’ takes place in the cue/delay period in both tasks, with and without the presence of the cue, hence the rise in rule sensitivity in frontal neurons. It appears that PPC neurons only participated this process in the presence of the cue.

### Ketamine has double-dissociable effects on spike-field consistency in the lPFC and PPC

While both neurons and LFPs in the frontoparietal network encode task-related information, it remains unclear whether the LFPs entrain task-related neuronal activities similarly in both areas, and how this may be affected by ketamine. Given that ketamine reduced frontoparietal connectivity, we expected that spike-phase coupling would also be reduced. Using cue/delay period activities, we calculated mean resultant length (MRL), which is a measurement of spike-LFP phase consistency [45]. To control for the confounding effect of spike counts, a bootstrapping approach was used. Additionally, we included only those neuron-LFP pairs that displayed significant spike-phase correlation against an empirical distribution constructed from shuffled spike times (see Materials and Methods).

When all frequency bands from theta (3-8Hz) to high gamma (50-80Hz) were considered, spike-phase consistency was higher in the PPC than the lPFC during the RV task (factorial ANOVA, area x task interaction: *F*_1,1634_ = 5.85, p = 0.016, post hoc Tukey’s test: p = 0.0056; Figure 8A) and was not different across areas in RM trials (p > 0.99). Among the frequency bands, spike-high beta (20-30Hz) phase consistency in the lPFC was significantly greater in the RM than in the RV task (p = 0.02). An exploratory analysis for the high-beta band revealed that this effect was carried exclusively by the antisaccade trials (area x task x rule interaction: *F*_1,248_ = 4.96, p = 0.027, antisaccades: p = 0.0045, prosaccades: 0.81; Figure 8B, solid gray vs black lines). This effect was not found in the PPC under either rule (Figure 8B, dashed lines). Thus, spike-field consistency modulated by the task in an area-dependent fashion in the frontoparietal network.

**Fig 8.**
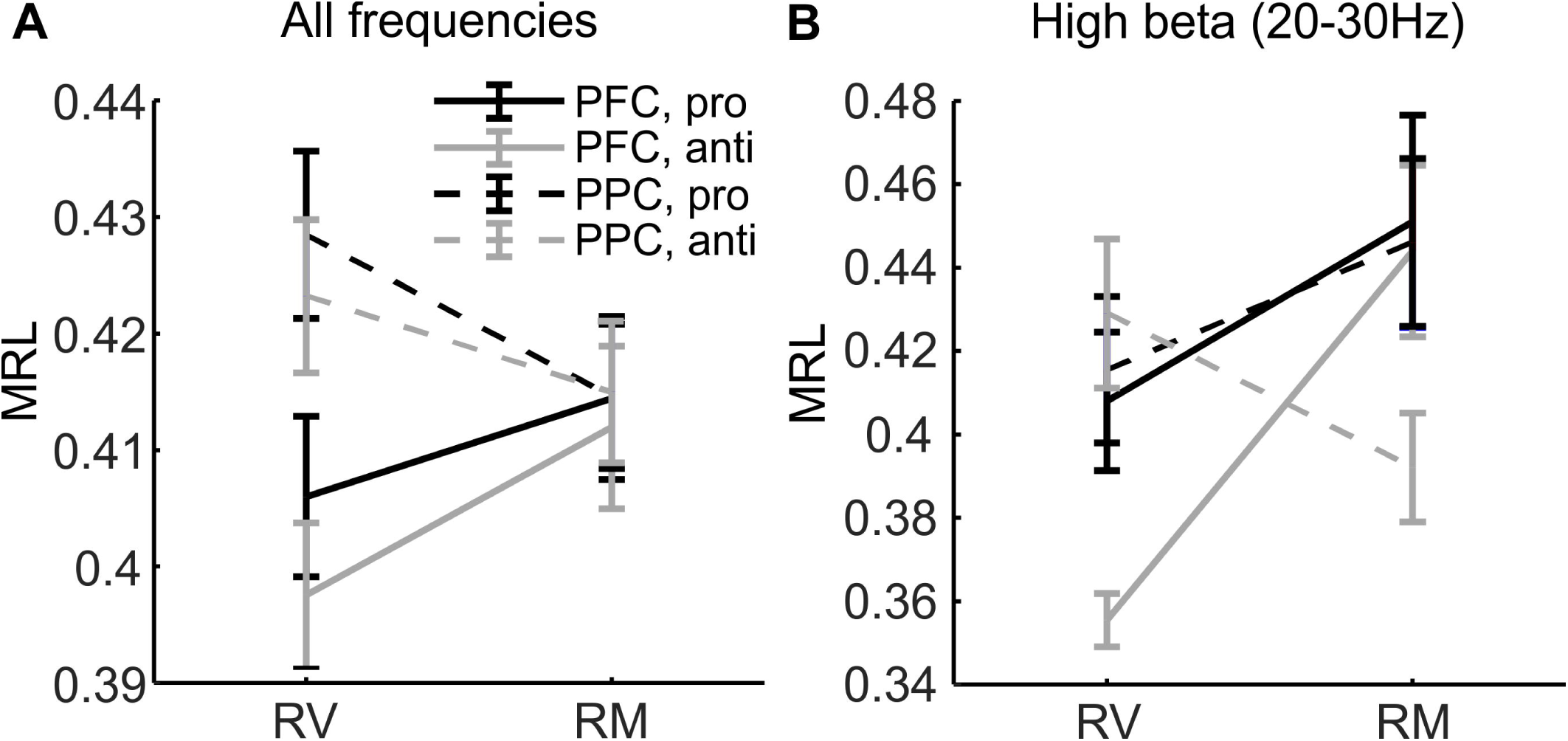
Spike-field consistency, quantified as the mean resultant length (MRL), depended on both the task and the brain region *before* ketamine administration. **A).** Averaged across all frequency bands, in visually-guide trials, MRL was stronger in the PPC (dashed lines); in memory-guided trials, MRL was equally strong across PFC (solid lines) and PPC. **B).** The low MRL in the PFC during the visual task was especially evident in antisaccade trials in the high-beta band. Solid lines: PFC. Dashed lines: PPC. Prosaccade trials: black, antisaccade trials: gray. Error bars indicate the standard deviation of the mean.

After establishing the task-relevance of spike-field consistency, we asked if ketamine affected this relationship. When all frequencies were considered, ketamine had double-dissociable effects on spike-field consistency in the two areas: it reduced MRL in the lPFC during RM but not RV trials (task x drug interaction: *F*_1,1564_ = 5.46, p = 0.02, post hoc test: RV: p > 0.99; STM: p = 0.019; Figure 9A, solid vs dashed lines). By contrast, it reduced MRL in the PPC during the RV but not RM trials (*F*_1,1857_ = 8.02, p = 0.0047, post hoc test: RV: p = 8.6 x 10^-6^; RM: p = 0.43, Figure 9B, solid vs dashed lines).

**Figure 9.**
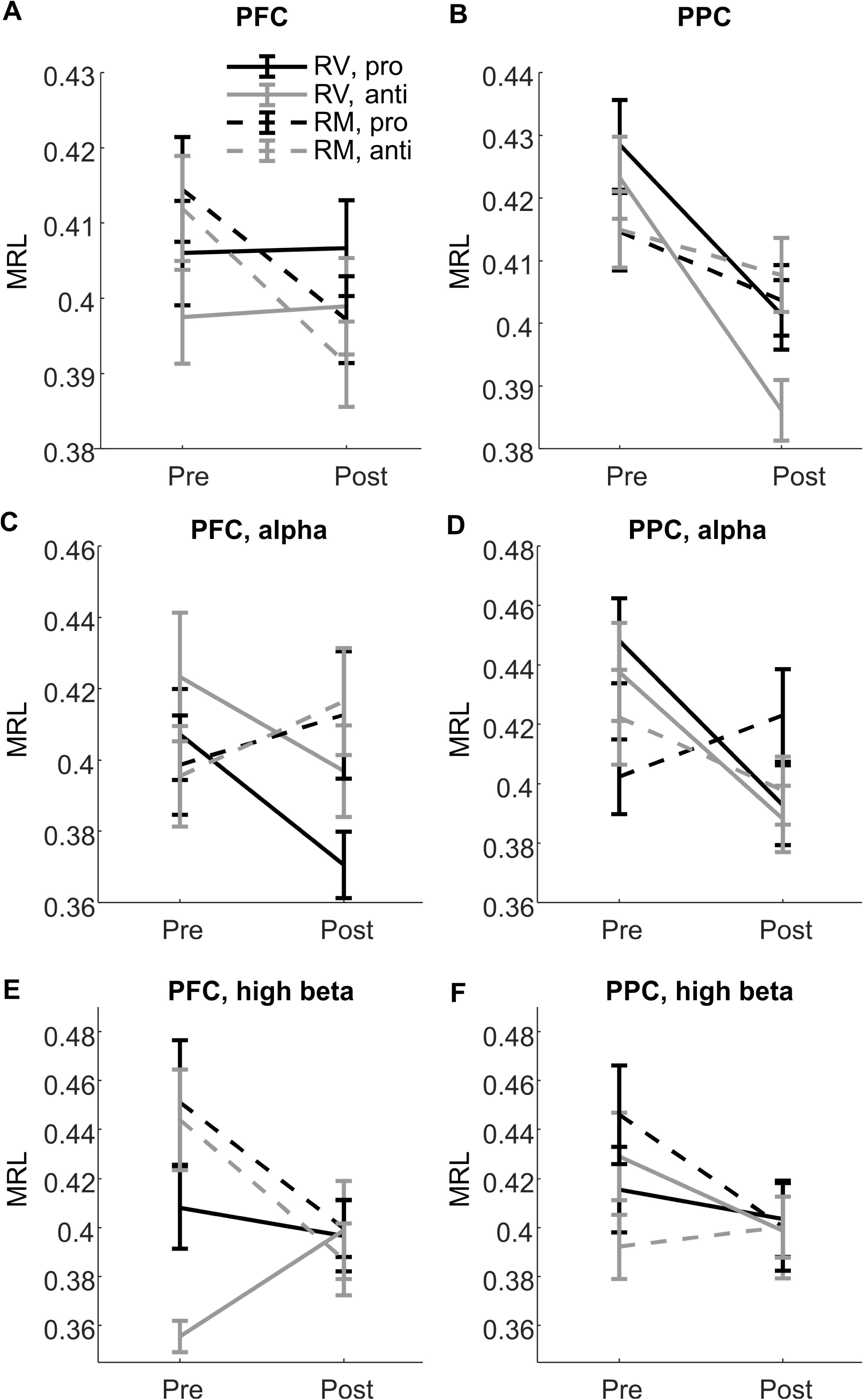
Ketamine had double-dissociable effects on spike-field consistency (MRL) in RV and RM tasks in lPFC and PPC. When all frequencies were considered, **A)** in lPFC, MRL was unaffected in RV trials (solid lines) but dropped significantly in RMtrials (dashed lines) following ketamine injections; **B)** in PPC, ketamine reduced MRL in RV (solid lines) but not RM task (dashed lines). Such effects were most strongly reflected at different frequencies: in the alpha band, **C)** ketamine did not significantly affect MRL in the lPFC, but **D)** reduced MRL in RV trials in the PPC. By contrast in high-beta band, E) ketamine significantly reduced MRL in the lPFC in RM trials, but F) did not affect the MRL in the PPC. Error bars indicate the standard deviation of the mean.

Additional post hoc tests showed that the effects in lPFC were mostly carried by the high beta band (20-20Hz), whereas the effects in PPC were strongest in the alpha band (8-13Hz). We therefore conducted additional analysis focusing on these two rhythms. Indeed, ketamine reduced the spike-alpha phase MRL in the PPC in RV (p = 0.0081; Figure 9D, solid lines) but not RM trials (p > 0.99, dashed lines), and reduced spike-high beta phase MRL in the lPFC in RM (p = 0.040; Figure 9E, dashed lines) but not RV trials (p = 0.98, solid lines). These effects were not found in the opposite brain area (Figure 9C and F).

Taken together, our analysis indicated a difference in how the lPFC and PPC contribute to working memory with or without a short term memory component, with PPC contributing more to the former and lPFC more to the latter. Moreover, the deleterious effects of ketamine tended to be stronger in an area when it wasthe most critical for task at hand..

Field potentials are known to partially reflect the summation of synaptic currents [46], which provide input to local neurons, whereas spikes reflect the output going to other regions. Since the lPFC and PPC are reciprocally connected, lPFC-PPC spike-field consistency could reflect directional connectivity from the frontal to parietal cortex in the ‘top-down’ direction, whereas PPC-lPFC spike-field consistency reflect connectivity from the parietal to frontal cortex in the ‘bottom-up’ direction. We therefore quantified these relationships using MRL and included only spike-field pairs that reached significance against an empirical distribution. We found that ketamine significantly reduced lPFC-PPC spike-field consistency during the delay period in the RM but not RV trials (factorial ANOVA, *F*_1,2307_ = 6.03, p = 0.014, post hoc test: RV: p = 0.91, RM: p = 0.01; Figure 10A), and reduced PPC-lPFC spike-field consistency in both trial types (*F*_1,2065_ = 53.91, p = 3.0 x 10^-13^, RV: p = 7.7 x 10^-6^, RM: p = 6.9 x 10^-4^; Figure 10B). This analysis demonstrated a deleterious effect of ketamine on frontoparietal synaptic communication in both directions, which is consistent with its negative impact on bidirectional Granger prediction.

**Fig 10.**
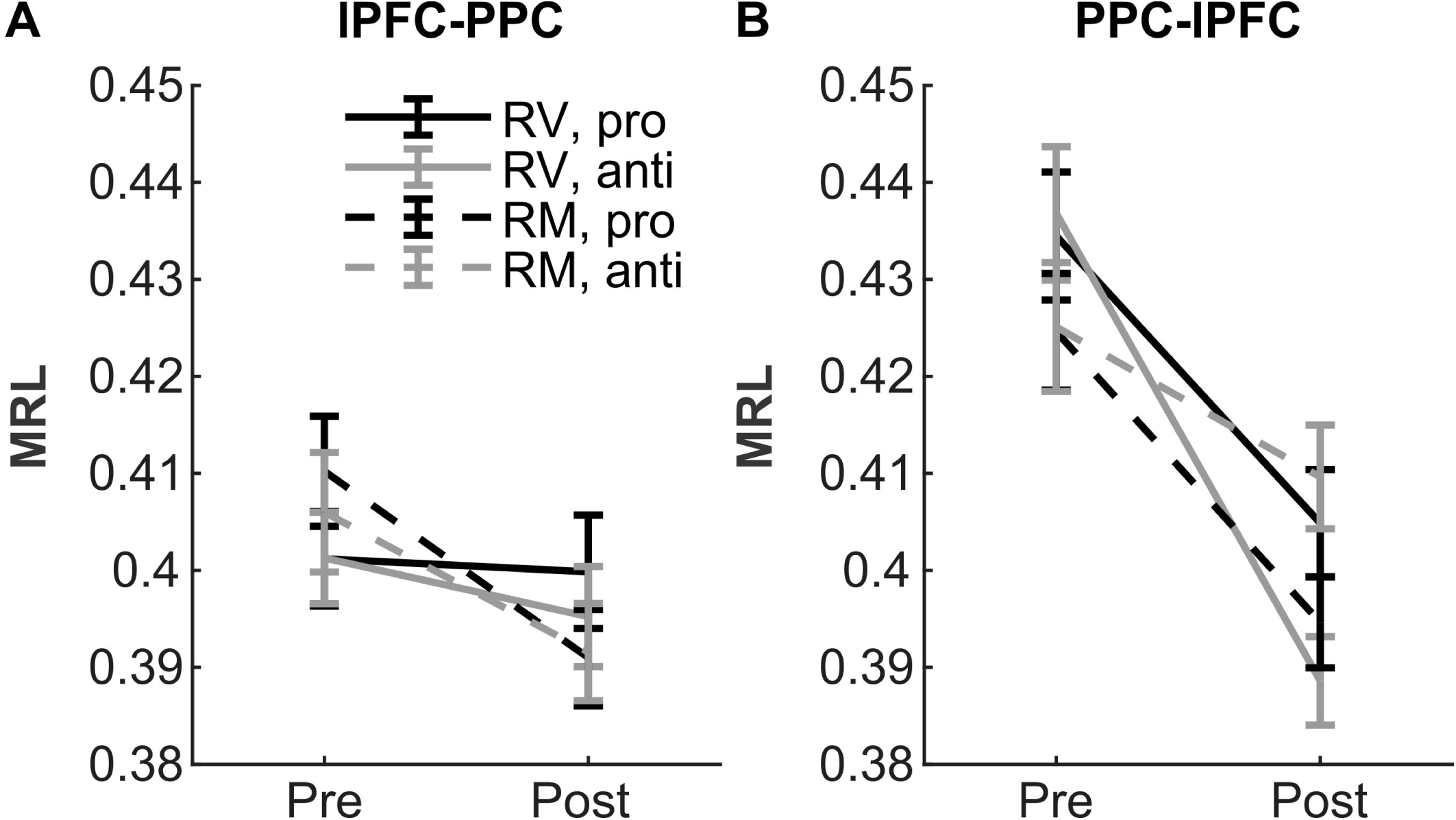
lPFC spike-PPC field consistency (**A**) and PPC spike-lPFC field consistency (**B**) estimated using mean resultant length (MRL). **A)** Ketamine reduced lPFC-PPC spike-field consistency in RM (dashed lines) but not RV trials (solid lines). **B)** Ketamine reduced PPC-lPFC spike-field consistency in both trial types. Error bars indicate the standard deviation of the mean.

## DISCUSSION

Through an analysis of single unit activity and local field potentials in macaque monkeys, we replicated the functional dysconnectivity in the frontoparietal network during working memory performance associated with schizophrenia [38–40]. We also simulated the reduction in low-frequency oscillatory power associated with this disorder [47] and with healthy volunteers receiving NMDAR antagonists [48–51]. This was accompanied by a task-dependent weakening in rule coding among frontoparietal neurons during the extended cue and delay period. Lastly, as a potential mechanism underlying ketamine-induced working-memory impairment, we found a task-dependent reduction in interareal spike-field correlations in the frontoparietal network.

### Acute ketamine administration has general effects on frontal and parietal activity

As previously reported for the lPFC [17,20] as well as in the current study, we observed an increased level of activity in the PPC and suggest the cause to be a general effect of ketamine on the cerebral cortex. First, ketamine is known to enhance spiking activity by increasing glutamate transmission through AMPA/kainate receptor currents [52,53]. Second, it may also exert a stronger NMDA receptor-mediated inhibitory effect on fast-spiking parvalbumin-positive (PV^+^) interneurons, thereby disinhibiting excitatory neurons [54–56]. Third, a metabolite of ketamine, (2S,6S;2R,6R)-hydroxynorketamine, stimulates AMPA receptors [57] and can further enhance the level of activity in cortical neurons regardless of their type. Future studies will verify if this effect may apply to sensory and motor cortical areas and to subcortical structures.

In oscillatory activities, ketamine weakened alpha-and low beta-band power in the PPC as it did in the lPFC [21], during both task and non-task periods. Similarly, this reduction in beta power has been observed in schizophrenia patients [48–51]. Beta-band oscillation has long been suggested to reflect long-range corticocortical communications [58]. Since cortical afferents often synapse with apical dendrites, which have a high density of NMDA receptors [59,60], NMDAR blockade is expected to disrupt frontoparietal functional connectivity [61]. This will be discussed in following sections.

Unlike our previous finding in lPFC [21], the enhancement in gamma rhythm (30-100Hz) was only observed during inter-trial intervals in the PPC but did not survive permutation test with family-wise error correction during other task periods. Despite of the lack of significance, the effect of ketamine on gamma is quite distinct from alpha/low-beta activity (Figure 3). Enhanced gamma activity has been observed in both EEG [47,62–64] and MEG [65] studies with schizophrenia patients. A similar observation was also made in healthy people receiving an acute, low dose of NMDA antagonists [66,67]. Previous studies in rodents suggested that ketamine may enhance gamma rhythm through GluN2A subunit-containing NMDA receptors in PV^+^ interneurons [68,69]. Although gamma-band oscillation in the cortex can be orchestrated by PV^+^ interneurons [70–72], the rhythmogenesis of gamma is highly diverse [73]. Here, we defined gamma rhythms more broadly than previous studies [74,75], and have not quantified the rhythmicity of gamma. Future studies combining data analysis and simulation will help reveal how ketamine affects cortical gamma from diverse sources.

Since LFPs are mainly a summation of synaptic currents [46], i.e. the input received by an area, spike-field consistency could reflect the level of coordination between input and output in that area. Intriguingly, we found a double dissociation in this measure: spike-field consistency was reduced by ketamine in the RM task in lPFC but not PPC, and in the RV task in the PPC but not lPFC (Figure 9). This finding indicates that the areas play a greater role in different tasks, which is consistent with our finding in rule coding by single units, discussed below.

### Acute ketamine administration impaired rule coding among frontoparietal neurons

Consistent with our previous studies involving lPFC alone [17,20], we observed a deleterious effect of ketamine on rule coding in both frontal and parietal neurons. In the RV task, while the cue remained visible, rule representation in the frontoparietal neurons was enhanced in the late phase of the cue period (Figure 7). This enhanced rule coding could be correlated with the retrieval of the pro/anti-saccadic rule to guide action selection. That is, a memory process was engaged even though short-term memory for the cue was not required.

Several studies have demonstrated that both lPFC and PPC contribute to short-term working memory by enhancing activity during the delay period [22,24,29,30,76]. Meanwhile, the two areas differ in anatomical and functional organization [34,77,78], and their roles in working memory [79]. Here, we observed an enhancement of rule encoding in the lPFC across tasks, whether the cue remained visible or not, and in the PPC only when the cue remained visible (RV task). This is consistent with previous findings that neural activity in the PPC is less generalized across tasks compared to the lPFC [28,80]. Alternatively, the lack of enhanced rule encoding at the population level may be related to the susceptibility of PPC neurons to distraction [22–25,27,81]. However, this does not mean that the PPC was not engaged by the short-term memory-guided task, during which intra-area coherence increased, as it did in the visually guided task (Figure 4). This task-related synchrony could be either intrinsic or driven by communication from the lPFC, as we observed task-related lPFC-PPC directional connectivity in the theta and high gamma band (Figure 5). Our findings support the idea that working memory in the intact brain is supported by cortical networks that include the lPFC and PPC [82–85]. We also showed that ketamine impaired rule-related memory processes through widespread effects on the frontoparietal network [41]. It remains to be tested whether the effect of ketamine extends to a greater cognitive control network involving additional cortical and subcortical areas in the primate brain [86,87].

### Acute ketamine administration led to task-related frontoparietal dysconnectivity

Given the profound effect of ketamine on glutamate transmission, it is not surprising that it also affected corticocortical dynamics. During off-task periods, we found that ketamine reduced alpha-band coherence between lPFC and PPC, and reduced theta coherence within each area. Consistently, reductions in the same periods were observed in lPFC-PPC and PPC-lPFC directional connectivity (Figure 6). Additionally, interareal spike-field consistency could reflect the extent to which output from area A in the form of spikes contributes to the synaptic input in area B. Therefore, we would expect this measure to show a similar effect of ketamine as did other measures of connectivity. Indeed, frontoparietal spike-field consistencies were reduced in a task-dependent manner (Figure 10). Resting-state functional MRI studies in patients with schizophrenia have reported either hypo-[38] or hyper-connectivity [37,88,89] in the frontoparietal network. Given that fluctuations in the BOLD signal occur at a much lower frequency than theta, it is difficult to compare our finding directly with this literature. Future electrophysiological studies using a longer delay period will serve to bridge findings from LFPs with the BOLD signal.

Compared to healthy controls, schizophrenia patients have lower frontoparietal connectivity during cognitive control tasks [36,39,40]. When given to healthy people, ketamine also disrupted task-related frontoparietal connectivity [90]. Furthermore, both a great need for cognitive control and better performance were correlated with enhanced frontoparietal connectivity [91,92]—a modulation that was impaired in patients [93]. Consistent with this literature, we found a task-related increase in theta, high beta, and high-gamma coherence between lPFC and PPC, which was attenuated by ketamine (Figure 5). Specifically, this attenuation was observed in both top-down and bottom-up directions, as well as the rule-related connectivity during the RM task (Figure 6). Finally, interareal spike-field consistency also dropped in a task-and direction-dependent manner (Figure 10).

Recently, Kummerfeld et al. (2020) studied the effect of NMDA receptor blockade via phencyclidine on frontoparietal dynamics during a cognitive control task. Using causal discovery analysis, they found that frontoparietal connectivity was weakened in the top-down, lPFC-PPC direction, but enhanced in the bottom-up, PPC-lPFC direction [41]. Two reasons may explain the discrepancy in our finding with PPC-lPFC directional connectivity: 1. it is difficult to establish a direct correspondence between Granger prediction and causal discovery analysis, and 2. Phencyclidine and ketamine have different effects on glutamatergic transmission: unlike ketamine, phencyclidine does not affect the firing rates of cortical neurons [94].

## Conclusion

Through a comprehensive analysis of both spiking and oscillatory activities in the frontoparietal network in macaque monkeys performing working memory tasks, we demonstrated the profound changes in network dynamics due to acute ketamine treatment. Our findings demonstrate the utility of the ketamine model in simulating a syndrome of functional dysconnectivity, and its potential for the exploration of novel treatment strategies for schizophrenia. Future theoretical work will reveal how NMDA receptor blockade and enhanced glutamate transmission lead to network dysconnection. Given the increasing popularity in the use of ketamine as an antidepressant [95], future studies are required to better characterize any long-lasting effect on the primate cognitive control network following repeated exposure to the drug.

## MATERIALS AND METHODS

### Animals

Two male rhesus monkeys (*Macaca mulatta*), weighing 8 kg (Monkey A) and 10 kg (Monkey C), were used in the study. As previously described (Johnston and Everling, 2006), animals were implanted with a plastic head restraint and trained to perform the task. Once trained, they were implanted with recording chambers over the posterior third of the prinicipal sulcus (lPFC chamber) and over the intraparietal sulcus (PPC chamber). Monkeys received postsurgical treatments including analgesics and antibiotics to minimize pain or discomfort, under the oversight of a university veterinarian. All proceduresperformed were approved by the Animal Care Committee of the University of Western Ontario Council on Animal Care and in accordance with the Canadian Council of Animal Care policy on laboratory animal use.

### Behavioral task

The task has been described in detail in previous papers from our group [17,20,21,42]. On each trial, animals were required to fixate a small white spot at the centre of the monitor display. After 500ms of fixation, a color cue replaced the white fixation spot and lasted for 400ms. On rule visible trials (RV), the cue presentation lasted for another 1000-ms period; on rule memorized trials (RM), the cue was extinguished after 400ms, followed by a 1000-ms delay with a small jitter, during which the screen remains dark (Fig. 1*A*). For Monkey A, a red cue indicated that a prosaccade was required for the subsequent post-delay peripheral stimulus, while a green cue indicated an antisaccade. For Monkey C, this colour contingency was reversed. The peripheral stimulus appeared 8° to either the left or right of the fixation spot at its offset. If the saccade landed within a window of 4° centred on the target location, the animal received a liquid reward. Trials involving prosaccades to the left and right and those involving antisaccades to the left and right were randomly interleaved. The animals’ eye positions were recorded and digitized at 1000Hz using an Eyelink 1000 infrared pupillary tracking system (SR Research, Mississauga, ON, Canada).

### Recording

At the start of a session, 4-6 tungsten recording electrodes were advanced through a grid placed over each brain region and removed afterwards. The locations of the electrodes were adjusted manually via screw microdrives and optimized for spiking activities within each session. Thefinal adjustment to any of the electrodes took place at least 20min before the onset of the recording session. Neural activities, including LFP and spike trains, as well as eye-tracking data were recorded using the PlexonIOmniplex system (Plexon, Dallas, TX, USA). Spiking activities were sorted manually using Offline Sorter (Plexon, Dallas, TX, USA). A portion of the local field potential data (LFPs) recorded from the lPFC of Monkey A have been used in the analysis of a previously published work[21].

### Drug administration

Each experimental session began with a pre-injection baseline period lasting at least 10min. The animals received a single intramuscular injection of either 0.4ml of ketamine or 0.4ml of 0.9% sterile saline. Ketamine at this dosage elicited cognitive deficits with minimal anesthetic effects in rhesus monkeys (Condy et al., 2005, Stoet and Snyder, 2006, Shen et al., 2010, Blackman et al., 2013). Both monkeys started at the dose of 0.4mg/Kg. Monkey A developed tolerance over time, whereas Monkey C was unable to perform the task at this dose. Hence, we analyzed the data after every session to confirm that 1) sufficient trials were performed, and 2) a significant change had been induced by ketamine in performance from 5 to 30min post injection compared to baseline. If this was not the case, the session was discarded, and the dose was increased/decreased by 0.1mg/kg for the next session. For Monkey A, the ideal dose was between 0.4—0.8mg/kg, and for Monkey C this was 0.1—0.15mg/kg. After the injections, the monkeys continued with the behavioral session for at least another 30min. Ketamine treatments were spaced by at least 72hrs to slow any development of tolerance.

### Preprocessing

From the 2 animals, a total of 323 channels had single unit activities and thus were included in the LFP analyses (lPFC and PPC: Monkey A: n = 109 and 110, Monkey C: n = 43 and 54). Because the behavioral effects of ketamine took no more than 5min to develop and returned to baseline 30min after injection, we compared the period from 5min to 30min after injection to the 10-min baseline preceding the injection. Single unit waveforms were manually sorted using Offline Sorter (Plexon, Dallas, TX, USA) and analyzed with custom software. LFP data were analyzed in MATLAB (MathWorks, Naticks, MA, USA, RRID:SCR_001622) using the FieldTrip toolbox (http://fieldtrip.fcdonders.nl/, RRID:SCR_001622) developed at the Donders Institute for Brain, Cognition and Behaviour (Oostenveld et al., 2011). The continuous signal was divided into discrete trials based on event timestamps. Trials in which delay-period LFP power exceeded 8 standard deviations from the average were excluded from the analysis. For the time-frequency presentation of LFP power, the data was tapered with a Hanning window with a size equivalent to 4 cycles of a given frequency, before fast Fourier transform was performed. To compare LFP between channels and animals and to reduce variability, we used z-score or decibel normalization for each channel at each frequency (Herrmann et al., 2014).

### Experimental design and statistical analysis

To quantify the drug effect accurately, we used a within-subject design, in which the animals’ neural activities as well as behavioral performance with ketamine on board were compared to the pre-injection baseline period within the same recording session. For behavioural performance and saccadic reaction times, we used mixed-model ANOVA, with time, rule and task as within-subject variables and drug as between-subject variables. For LFP data, we used repeated-measures ANOVA, with treatment (pre-vs post-injection), rule, and task as within-subject factors. The following subsections provide the details to each statistical analysis we performed.

#### Permutation test

Statistical testing on the time-frequency LFP power map was conducted using a nonparametric cluster-based method. Specifically, a map of *t*-statistics was calculated between the two conditions. The significance level was then determined from a distribution of *t*-statistics generated by 1000 iterations of pooling and random splitting of the data. Original *t*-statistics that were greater than 99% of the generated distribution were considered significant. Significant *t*-statistics were then clustered using the *clusterdata* function in MATLAB. The significant time-frequency pairs are demarcated with a black contour in Figure 3A and B.

#### De-biased weighted phase-lag index (dWPLI)

We used the dWPLI to quantify the phase coherence between LFP channels within and across brain regions. This estimate is based on the imaginary component of the coherency between LFP series and independent of LFP power, with additional improvements: compared to the phase locking value (PLV), the dWPLI is less affected by phase delays; and compared to the phase lag index (PLI) it is less affected by noise and has enhanced statistical power [43]. Lastly, the ‘de-biasing’ refinement removed the sample-size bias [43].

We used the FieldTrip toolbox (http://fieldtrip.fcdonders.nl/, RRID:SCR_001622) (Oostenveld et al., 2011) to first calculate the cross-spectra using the same windowing function as we did for power spectra (see *Preprocessing*). We then used the ft_connectivityanalysis.m and set the method to ‘wpli-debiased’ to calculate dWPLI for each time-frequency pair. To examine task-related coherence, we z-score normalized the delay-period dWPLI against the dWPLI during ITIs (Figure 4 and 5).

#### Granger prediction (GP)

To analyze the task-related and ketamine-induced effects on *directional* connectivity in the frontoparietal network, we calculated Granger prediction using the Multivariate Granger Causality (MVGC) toolbox [44]. For each pair of channels, the time-domain data from the delays and ITIs were separately fit with a vector autoregressive (VAR) model, then the VAR coefficient and residual covariance matrices were used to calculate the multi-variate Granger prediction (GP). We then compared the Granger prediction using repeated-measures ANOVA, with direction, task, and treatment as within-subject variables (Figure 6A). Additionally, we analyzed the task-specific GP, obtained by normalizing delay-against ITI-GP (Figure 6B).

#### Rule Selectivity (d’)

To quantify the rule information contained in the activities, we calculated the Rule Sensitivity, or absolute d’, for each neuron:

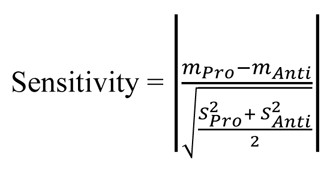

Where *m*_*Pro*_ and *m*_*Anti*_ denote its mean firing rate, and *S*^2^_*Pro*_ and *S*^2^_*Anti*_ denote its variance in firing rates during pro-and anti-saccade trials, respectively. We calculated this value for each epoch of interest: fixation, cue, delay (or the later part of the cue period in visually guided saccadic trials) and peri-saccade periods. We used a bootstrapping procedure to control for the effect of sample size on d’. Each calculation of the numerator and denominator of d’ was done with a randomly selected set of 15 pro-and anti-saccade trials, the process was repeated 1000 time and the results were averaged before the final absolute d’ was obtained (Figure 7A, B). To identify the neurons with a significant sensitivity to the rules, we built an empirical distribution for each neuron by shuffling and calculating the d’ using the selected set of pro-and anti-saccade trials 1000 times. A neuron was deemed significantly sensitive to the rules if the original absolute d’ was greater than 97.5 percentile of the empirical distribution (α = 0.05, two-tailed).

#### Spike-field phase consistency

We used the mean resultant length (MRL) to quantify the spike-field phase consistency (Sigurdsson et al., 2010). For each neuron, a spike is fired at a unique phase with the LFP oscillation at each frequency. This phase can be represented by a unit vector with length equal to one. The MRL for a given frequency is defined as the vector sum of these unit vectors representing the phases when the spikes occurred, divided by the number of spikes—or number of unit vectors—included in the calculation, resulting in a number between 0 and 1. Thus the MRL is independent of LFP power; it is larger when these phases are similar than when they are dissimilar from spike to spike. However, the MRL is negatively correlated with the sample size of spikes: the more spikes a neuron fired, the greater the variance in the spike-associated oscillatory phases tends to be, resulting in a smaller vector sum and smaller MRLs. We controlled for the sample size bias by bootstrapping: for each neuron, eight spikes were randomly selected to calculate a single MRL, then the process was repeated 1000 iterations and the results were averaged to obtain the final MRL [21].

